# A genomic prediction framework for growth traits in black soldier fly (*Hermetia illucens*) breeding programmes

**DOI:** 10.64898/2026.11.05.764768

**Authors:** Kishor B. Gowda, Cecile Massault, Dean R. Jerry, Kyall R. Zenger

## Abstract

**Background:** Genomic prediction has been applied in livestock, plant and aquaculture species to accelerate genetic gain. However, its application and potential still remain unexplored in farmed black soldier fly (BSF), an emerging globally important insect production species. Due to the diversity of BSF feed sources, selection strategies need to be evaluated across various feedstocks and production environments to aid in building genomic selection into BSF breeding programmes. In this study, 2,083 BSF larvae from the base population were reared on three diets (SYK: Soy-okara, BSG: brewers’ spent grain and FVW: fruit and vegetable waste), and 536 from the F7 population were reared on a chicken starter feed diet. Larvae were phenotyped for four growth traits (LBW, larval body weight; LL, larval length; LW, larval width; LSA, larval surface area) and genotyped across 4,680 SNPs. Genomic prediction was evaluated using seven methods under additive (A) and additive-dominance (AD) models, considering across-and within-diet effects, genotype by diet (G × D) interactions, differences amongst reference and test generations, and SNP densities.

**Results:** Performance varied across all scenarios, with LBW showing higher predictive ability (PA) among the growth traits. Accounting for dominance generally improved PA. Multivariate models accounting for G × D performed similarly to, or marginally better than, within-diet predictions. Cross-diet prediction was more variable and generally lower than within-diet prediction, showing reduced transferability across untested nutritional environments. Reducing the SNP density from 4,680 to 3,000 SNPs maintained comparable PA; however, PA decreased below 3,000 SNPs. Across multiple generations, predictive performance declined, resulting from reduced genetic connectivity and possible genotype-by-environment (G × E) interactions between the reference and testing populations.

**Conclusions:** Overall, these results provide an application-based framework for genomic prediction in BSF in semi-or commercial production settings. Results also highlight the importance of the genetic model, representative reference populations, and appropriate marker densities for genomic selection.

## Background

The insect farming industry has grown rapidly in recent decades, with the market projected to reach USD 9.8 billion by 2030 [1]. Insect farming has applications in food and feed production, similar to conventional animal and agricultural production systems. Insect-derived ingredients are used in animal and aquaculture feed as alternative protein and fat sources, and edible insects are used for human consumption [2]. As the industry matures, it is shifting from traditional mass-rearing systems to more systematic, efficient production through automation, streamlined processes, and enhanced sustainability [3]. As these systems become more intensive, companies are moving toward implementing genetic improvement programmes, including cross-breeding and selective breeding to enhance productivity by improving important economic traits such as growth, health and production efficiency [2, 4]. These programmes are necessary to manage and improve the genetic merit of the breeding populations.

The Black soldier fly (*Hermetia illucens*, Diptera: Stratiomyidae; BSF) has emerged as an important species in the insect production industry, supporting the circular economy by converting organic waste into useful biomass. Selective breeding programmes are implemented to improve production efficiency and enable efficient bioconversion in BSF production systems [5–7]. However, this requires reliable estimation of the genetic merit of individuals/lines within the breeding programme. BSF larvae growth and production traits vary substantially, and studies have estimated genetic parameters such as heritabilities and correlations across different genetic strains and production environments [8, 9]. Recent developments in genomic resources for BSF, genome assembly and resequencing [10, 11] and genotyping tools such as SNP arrays and genotyping by sequencing (GBS) [12, 13] have enabled characterisation of genetic and population structure worldwide. However, to date, no published studies have validated genomic information to predict genetic merit, or how effectively it can be used to predict genetic merit in BSF breeding colonies/programmes. Therefore, the genomic prediction reliability, accuracy, and transferability need to be studied and quantified.

Modern selective breeding programmes for industrialised plant and animal sectors use genomic selection, where an individual’s genetic merit is estimated as genomic estimated breeding values (GEBVs) using genome-wide molecular markers and selected based on these estimates [14]. With advances in genomic technologies and reduced costs, genomic prediction is becoming more feasible in livestock, plants, and aquaculture [15–17]. Genomic prediction models applied in livestock and aquaculture commonly focus on the additive genetic component and ignore non-additive components, such as dominance. Studies in pigs and sheep reported improved prediction performance when considering dominance along with additive effects compared with additive-only models [18, 19]. However, the advantage of including dominance may depend on the population and trait architecture [20]. This consideration needs evaluation, particularly in colony-based and mass-rearing insect species like BSF. Captive breeding may change allele frequencies, heterozygosity, and genetic relationships because of founder effects, genetic drift, inbreeding, and selection pressure, potentially affecting the contribution of dominance to genetic variance [21, 22]. Additionally, breeders often cross different genetic strains to increase genetic diversity and potentially exploit hybrid vigour driven by dominance effects [23]. Therefore, comparing additive-only and additive-plus-dominance models requires validating and evaluating genomic prediction accuracy.

BSF larvae (BSFL) are often raised on diverse organic substrates/diets that vary in nutritional composition and provide different production environments. Studies have indicated genotype-by-diet interactions (G × D) in BSF populations, as genotype expression depends on diet and genotypes re-ranking as they move from one diet to another [9, 24, 25]. These interactions may require developing diet-specific strains or generalist strains that perform well across diets. G × D interactions may reduce the transferability of genomic predictions between diets if the prediction model does not account for these interactions. Therefore, a multivariate genomic prediction model is a framework that accounts for these interactions through genetic correlations of the same trait across diets [26]. These approaches have been tested in various animal, plant, and aquaculture breeding programmes [27, 28] and help evaluate within-diet and across-diet prediction performance. This model structure can also be used to evaluate cross-environment/diet prediction, where training is done on known environments, and performance is predicted and evaluated in a new environment.

The genomic prediction accuracy may plateau after a certain number of markers, even as marker density increases, depending on the trait genetic architecture and how well the available markers capture underlying genetic variability [29]. Recent aquaculture studies indicate that moderate marker densities are sufficient to achieve reliable prediction accuracy; therefore, adding extra markers may add little gain but increase genotyping and computation costs [30, 31]. Evaluating the effect of marker density on prediction accuracy and identifying the optimal marker density to achieve reliable prediction accuracy will help streamline genomic evaluation methods and reduce genotyping costs. Additionally, evaluating the prediction ability across generations is necessary to evaluate temporal transferability in breeding programmes [32]. Forward prediction assesses prediction performance in future generations using current-generation information and supports future selection decisions. Backward prediction, by contrast, assesses previous-generation prediction performance based on current-generation information and overlooks temporal transferability. This may be relevant for BSF breeding programmes, which require introducing genetically diverse germplasm into the breeding nucleus to maintain diversity and alleviate domestication bottlenecks. Backward prediction may provide a baseline for how well current information can be used to evaluate previously established or archived germplasm.

This study evaluates the performance of different genomic prediction methods for BSF growth traits, including genomic best linear unbiased prediction (GBLUP), Bayesian, and machine learning approaches. In particular, the study compares additive and additive-dominance models, accounts for G × D interactions and cross-diet prediction and evaluates the effect of different marker densities on prediction performance. Additionally, it assesses the transferability of genomic prediction across multiple generations through forward and backward prediction scenarios.

## Methods

### BSF population and phenotypes

The BSF populations used in the current study consist of two distinct populations established from the same original genetic background: (1) the base population, which corresponds to the dataset described in [9] and (2) the F7 population representing the 7^th^ generation of the BSF colony maintained at JCU, Australia.

### Base population

Briefly, the base population was established by collecting BSFL (n∼1,000) from ten wild BSF populations from across the species distribution in Australia and mixing them in a mass-rearing breeding design. The mixed population was maintained and mixed in the same conditions for three further generations. In the 4^th^ generation, 53 egg clutches were collected from this mass-rearing colony on a single day, allowed to hatch in the unified nursery, and reared on three experimental diets: Soy-okara (SYK), brewers’ spent grain (BSG) and fruit and vegetable waste (FVW) in triplicate trays. Phenotypic measurements were collected from 2,097 BSFL, consisting of larval body weight (LBW), larval length (LL), larval width (LW), and larval surface area (LSA), and were recorded from fifth-instar larvae on days 13, 14, and 15 after egg collection. Subsequently, tissue samples were preserved for genotyping.

Phenotypes (LBW, LL, LW and LSA) of the base population were adjusted using the fixed-effect models in *ASReml-R v4.2* [33]. For the overall analysis, phenotypic records of larvae reared across diets were considered, with colour (RGB values), a potential indicator of physiological or developmental differences among larvae, fitted as a covariate, while diet, sampling day and tray nested within diet were fixed effects. For the diet-specific analysis, colour (RGB values) was fitted as a covariate, while tray, sampling day, and tray nested within sampling day were fixed effects. The adjusted phenotypes are used as response variables in the genomic prediction models (Section 2.5).

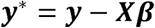

Where,Y* is the adjusted phenotype *β*, is the vector of fixed effects and *X* is the design matrix relating the fixed effects to the observed phenotype *y*.

The adjusted phenotypes were checked for normality and were near-normally distributed (Additional file 1, Figures S1 and S2).

### F7 population

The F7 population was sampled from the 7^th^ generation of the ongoing BSF breeding colony maintained at JCU. This colony originated from the same genetic background as the base population described in [9]. In each generation, at least 300 egg clutches/families were collected, and all other husbandry practices were similar to the base population breeding design. In the 7^th^ generation, approximately 30 egg clutches were collected on a single day from 70 mating pairs separated during mating. Offspring hatched in a common nursery environment and were reared on a commercial chicken starter feed (CSF) diet (15.5% crude protein) in multiple trays. In total, 576 BSFL were phenotyped at the fifth instar for LBW (mean ± SD: 150.97 ± 29.36 mg) and LSA (mean ± SD: 64.51 ± 12.07 mm^2^), and tissue samples were collected and preserved for subsequent genotyping. All mating procedures, nursery management, larval rearing conditions, and phenotyping protocols followed the methods described in [9].

F7 population phenotypes (LBW and LSA) were adjusted with colour as a covariate and tray as a fixed effect, as previously described.

### Genotyping, quality control and population structure

The preserved tissue samples were used for gDNA extraction with the Chemagic™ 360 magnetic bead extraction system (PerkinElmer Inc.) following the manufacturer’s protocol for the CMG-723 tissue kit. Following quality control, samples were genotyped using a custom-designed 6K Allegro targeted SNP panel developed for BSF [12] at AGRF Ltd, Brisbane, Australia. In total, 2,097 individuals from the base population dataset and 576 individuals from the F7 population dataset were genotyped. Genotype quality control was performed using *PLINK v1.9* [34] based on an individual call rate >90%, a SNP call rate >90%, and a minor allele frequency (MAF) >0.5%. After quality control, 4,680 SNPs were retained across 2,083 individuals in the base population and 536 individuals in the F7 population.

The population structure of the base and F7 populations was analysed using the first and second principal components (PCs) calculated using the *--pca* option in *PLINK v1.9* [34], based on the filtered dataset comprising 4,680 SNPs across 2,619 individuals from both populations. Population structure was visualised in R using *ggplot2* [35] package.

### Genomic prediction models

Genomic predictions were conducted to evaluate the performance of different statistical models for genomic estimated breeding value (GEBV) prediction of BSFL growth traits (LBW, LL, LW, and LSA). The models evaluated were GBLUP, Bayes A (BA), Bayes Bπ (BBπ), Bayes Cπ (BCπ), Bayes LASSO (BL), Bayes R (BR) and Random Forest (RF). Models were fitted using either additive (A) or additive plus dominance (AD) genetic effects.

### GBLUP

GBLUP was implemented in *ASReml-R v4.2* [33] using the residual maximum likelihood (REML) approach, with additive (A) and additive-dominance (AD) models.

a) Additive model (A)

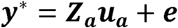

b) Additive-dominance model (AD)

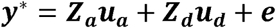

Where *Y*\* is the vector of adjusted phenotypes. The terms *u_a_* and *u_b_* represent vectors of random additive genetic and dominance effects, respectively, and *e* represents the vector of random residual effects. The matrices *Z_a_* and *Z_b_* are the incidence matrices linking observations to additive genetic and dominance effects, respectively.

The random effects were modelled as normally distributed, with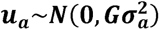, where 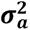 is the additive genomic variance, and *G* is the genomic relationship matrix (GRM) constructed using [36] method in *snpReady* [37] package in R. Similarly, 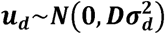 , where 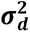 is the dominance genomic variance, and *D* is the dominance relationship matrix (DRM) constructed using [38] method in *AGHmatrix* [39] package in R. Residual effects were assumed to follow 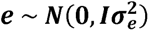, where 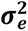 is the residual variance, and *I* is the identity matrix.

Multivariate A and AD models were used for genomic prediction across diets to account for G × D interactions. For each LBW and LSA, we treated adjusted phenotypes from SYK, BSG, and FVW as separate but correlated traits. Additive and dominance genomic relationships were modelled using the *G* and *D* matrices as described above, allowing unstructured genetic covariance structure between the diets.

Bayesian models

Bayesian models were fitted as marker effect models compared to the individual-level GBLUP models. A and AD models were considered;

a. A model

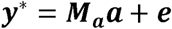

b) AD model

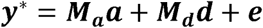

Where *M_a_* and *M_d_* are the genotype matrices for additive and dominance effects, respectively; *a* is the vector of random regression coefficients for additive marker effects, and *d* is the vector of random regression coefficients for dominance marker effects. The remaining notations are the same as those mentioned in GBLUP.

Bayesian models, including BA, BBπ, BCπ, and BL, were implemented using the *BGLR* package in R for the univariate analysis. Models were fitted using the prior distributions and marker effect specifications previously described in [40].

For the multivariate analysis across diets, we implemented a Bayesian spike-slab model in BGLR, using the same marker-effect specification as the BCπ model.

The BayesR model was implemented in the *hibayes* [41] package in R. BayesR assumes that markers’ effects arise from a mixture of distributions with different variances, allowing markers to contribute effects of different magnitudes.

All Bayesian analyses used 12,000 iterations, including a burn-in period of 2,000 iterations.

Random Forest

Random forest (RF) was implemented in the *ranger* [42] package in R for the univariate models, whereas for the multivariate models, it was implemented in the *randomForestSRC* [43] package in R. The model hyperparameters, including number of trees (150, 300), node sizes (5, 10), and number of candidate split variables 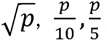 and 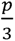 where *_p_* is the number of SNPs), were optimised within each training dataset. The best parameter combination was selected for each training dataset using nested 5-fold cross-validation.

### Genomic prediction scenarios and cross-validation

Genomic prediction models were investigated under multiple scenarios to assess trait definitions, diet-specific effects, G × D interactions, marker density, and forward and backward prediction. The following scenarios were tested:

1. Genomic prediction based on the base population dataset on individual growth traits, including LBW, LL, LW and LSA
2. Genomic prediction based on the within-diet (SYK, BSG and FVW) in the base population for the growth traits, including LBW and LSA, for each diet
3. Genomic prediction by accounting for the G × D interaction in the base population using the multivariate model by considering the trait in each diet as separate and correlated
4. Genomic prediction to evaluate cross-diet prediction in the base population, where training is on individuals from two diets and predicting the individuals in the 3^rd^ diet
5. Evaluating the different marker panel densities in the base population using the full 4,680 SNP panel and reduced panels containing 4,000, 3,000, 2,000, 1,000 and 500 SNPs; markers were randomly chosen by giving weight to the total markers on each chromosome (Additional file 1, Figure S3). In addition, the top 1,000 markers were selected based on GBLUP-derived SNP effects using the method of [44]
6. Forward and backward prediction

Forward prediction: training on the base population and predicting the F7 population. Backward prediction: training on the F7 population and predicting the base population.

Prediction performance was evaluated using random 5-fold cross-validation (4 folds for training and 1 fold for validation), with 20 replications, except for scenarios 4 and 6, where training and validation populations were predefined.

### Prediction performance evaluation

The prediction performance of the models was assessed using three metrics: predictive ability (PA), mean squared error prediction (MSEP) and prediction scale bias.

Predictive ability (PA) was calculated as the Pearson correlation coefficient (r) between the adjusted phenotype 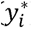 and genomic estimated breeding value (GEBVi)of the individuals in the validation set;

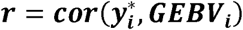

The mean squared error prediction (MSEP) was calculated using the following formula:

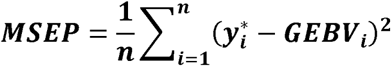

Where n is the total number of individuals in the validation set,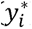 is the adjusted phenotype of the i^th^ individual, and GEBVi is the genomic estimated breeding value of the i^th^ individual.

The prediction scale bias (bi) was estimated as the slope of the linear regression of the adjusted phenotype on the genomic estimated breeding values in the validation set:

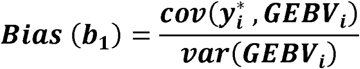

## Results

### Population structure

PCA was performed to evaluate the genetic structure of the base (2,083 individuals) and F7 (536 individuals) populations. PC1 and PC2 explained 18.66% and 13.31% of the total genomic variance, respectively. The base population showed two broad, separate clusters, whereas the F7 population comprised a single cluster. Individuals from the base population were distributed across all three diets, with no clear diet-specific clustering. The two populations did not overlap in the PCA plot, indicating genetic differentiation between the base and F7 populations despite their shared genetic background. This was supported by a genome-wide weighted *F_ST_* of 0.063 ± 0.004, indicating quantifiable allele-frequency differences between the base and F7 populations (Fig. 1).

**Fig. 1:**
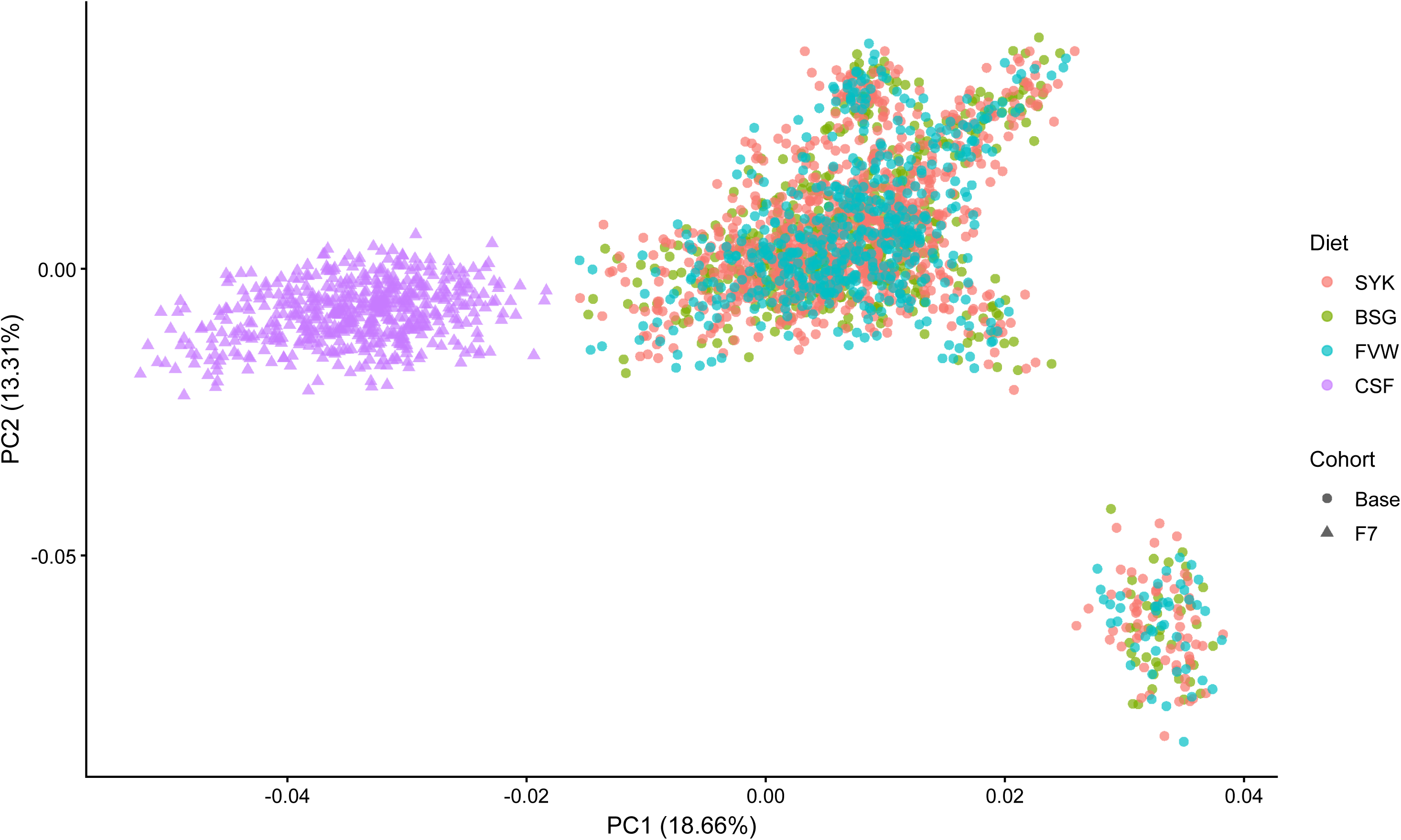
Principal component analysis (PCA) illustrating the distribution of the individuals across the diets (SYK: soy-okara, BSG: brewers’ spent grain, FVW: fruit and vegetable waste, CSF: chicken starter feed) in the base and F7 populations based on the first two principal components (PC1 and PC2).

### Genomic prediction performance across different validation scenarios

Effect of prediction method and model on BSFL growth traits

Prediction performance was measured using PA, which varied with method and across four BSFL growth traits (Fig. 2). The AD models generally performed better, with higher PA across all traits and methods than the A models. The mean PA was highest for LBW, ranging from 0.29 to 0.41 for A models and from 0.40 to 0.43 for AD models, followed by LSA, LL, and LW (Additional file 2, Table S1). The MSEP dropped across all the methods and traits when dominance was included compared to A models. Bias was close to unity across models, methods, and traits (Additional file 2, Table S1).

**Fig. 2:**
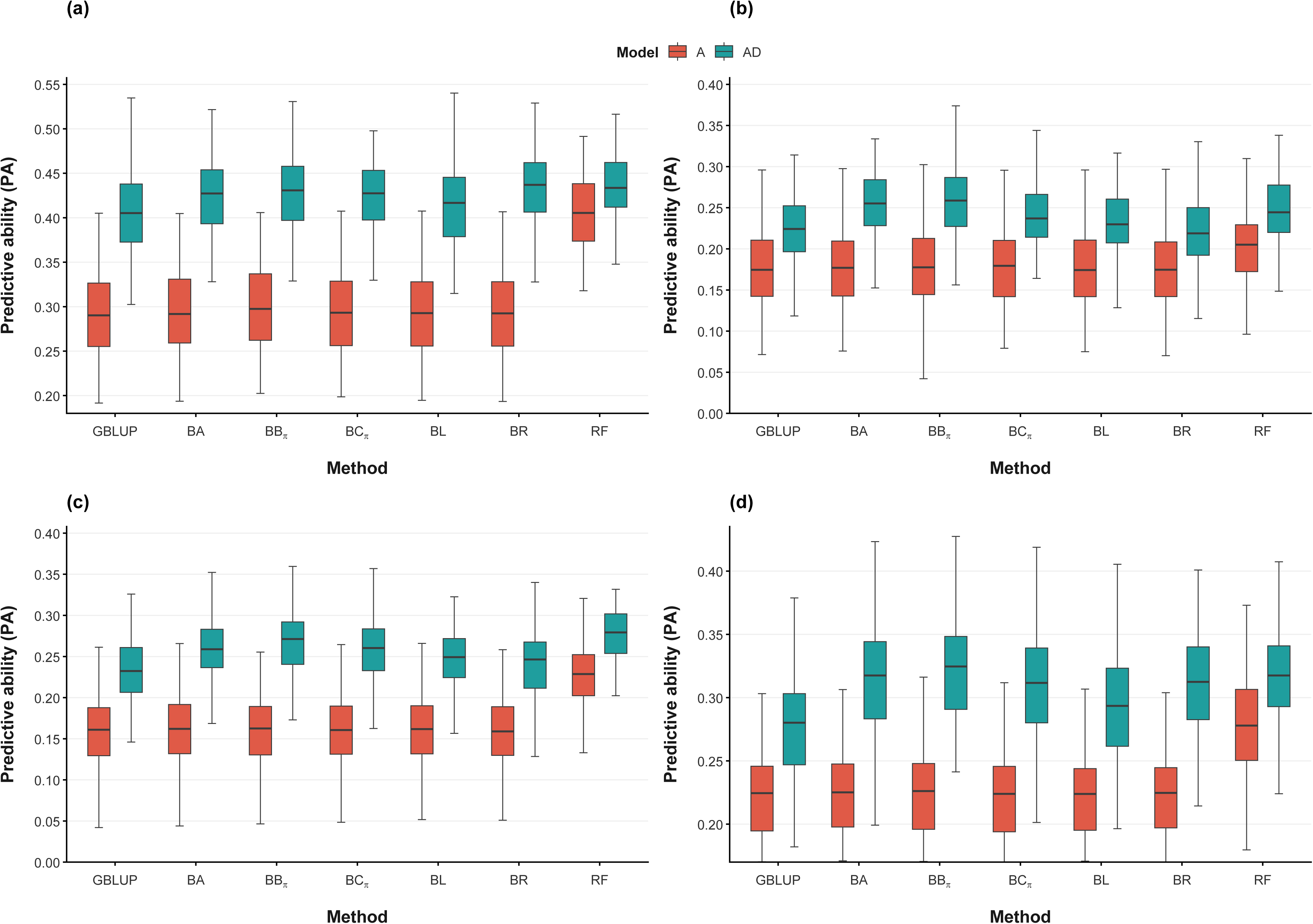
Genomic predictive ability (PA) of different methods: genomic best linear unbiased prediction (GBLUP), Bayes A (BA), Bayes Bπ (BBπ), Bayes Cπ (BCπ), Bayes LASSO (BL), Bayes R (BR), and random forest (RF), using additive (A) and additive-dominance (AD) models for four BSFL growth traits: (a) larval body weight (LBW), (b) larval length (LL), (c) larval width (LW) and (d) larval surface area (LSA)

Among the methods, RF generally performed better across all the traits and models with higher PA, particularly with A models. In comparison with GBLUP, RF PA was 41.4% higher for LBW, 16.7% for LL, 43.8% for LW and 45% for LSA. However, when the dominance effect was included in the AD models, these differences were reduced, with RF performing relatively better at about 10%, 13.6%, 16.7%, and 14.3% for LBW, LL, LW, and LSA, respectively, compared with GBLUP. Bayesian methods showed broadly similar PA to GBLUP across traits. Under AD models, RF performed slightly better than Bayesian methods for LBW and LSA.

### Effect of prediction method and model on within-diet growth traits

Prediction performance varied among the diets and between A and AD models within diets (Fig. 3). Across methods, SYK consistently showed the highest PA for both LBW and LSA, followed by BSG and FVW. For LBW, SYK PA ranged from 0.40 to 0.54 under the A model and from 0.54 to 0.59 under the AD model. For LSA, SYK PA ranged from 0.29 to 0.33 under the A model and from 0.36 to 0.40 under the AD model (Additional file 2, Table S2). In FVW, PA was similar between the A and AD models for both LBW and LSA; by contrast, SYK and BSG differed between the A and AD models. RF showed higher PA than other methods under A models across all diets for LBW, but only a slight improvement under AD models. Bayesian models looked similar under A models for all diets in LBW and LSA but varied in AD models; BA and BBπ performed better for LSA.

**Fig. 3:**
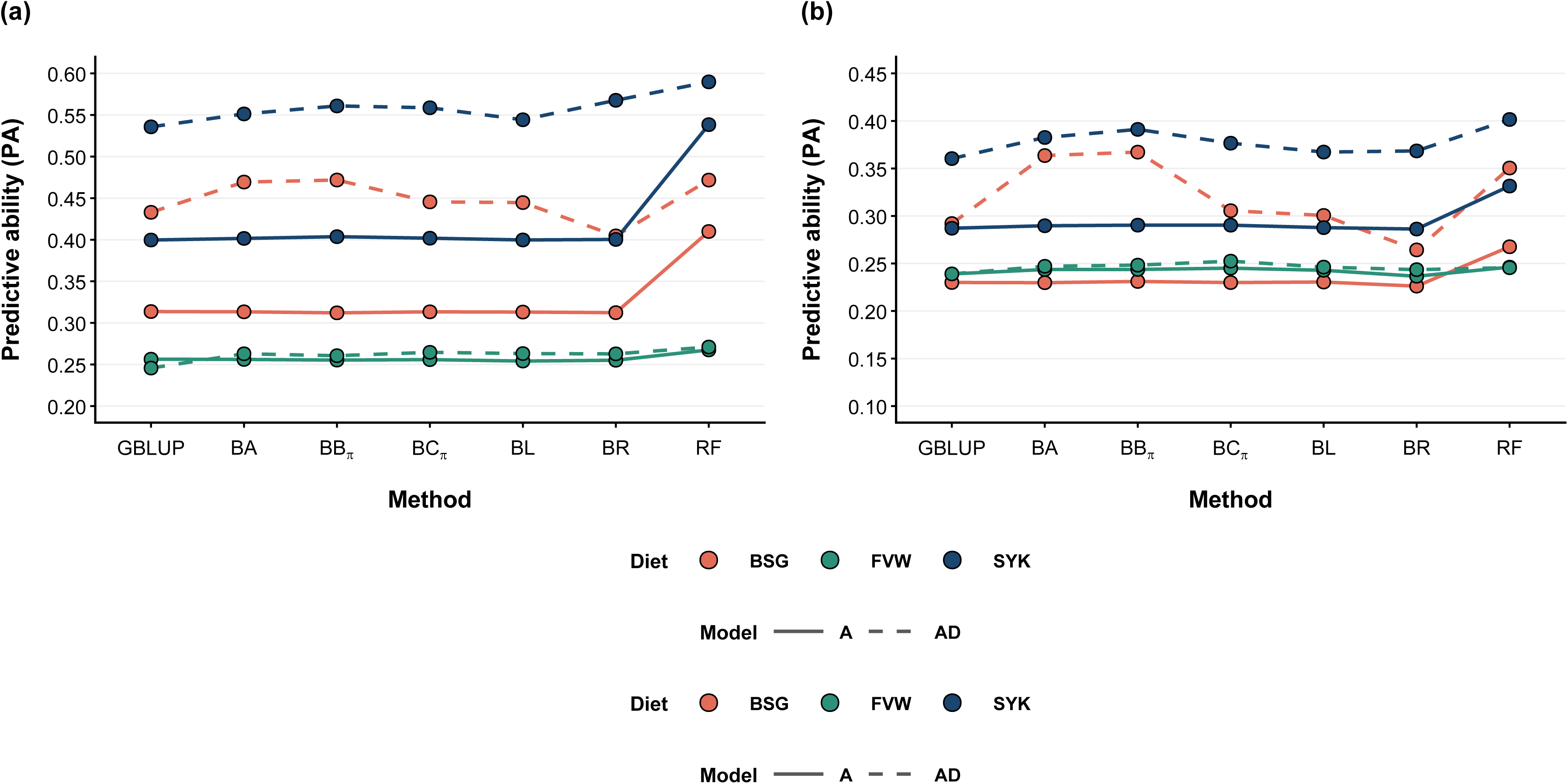
Genomic predictive ability (PA) of different methods: genomic best linear unbiased prediction (GBLUP), Bayes A (BA), Bayes Bπ (BBπ), Bayes Cπ (BCπ), Bayes LASSO (BL), Bayes R (BR), and random forest (RF), using additive (A) and additive-dominance (AD) models for within-diet (SYK: soy-okara, BSG: brewers’ spent grain, FVW: fruit and vegetable waste) prediction of BSFL growth traits: (a) larval body weight (LBW) and (b) larval surface area (LSA)

### Effect of accounting for G × D interactions on prediction performance

The multivariate analysis showed minor changes in PA compared with the univariate analysis across methods and models (Fig. 4). For BSG, PA under model A in RF improved from 0.41 to 0.47 for LBW and from 0.27 to 0.31for LSA. In contrast, multivariate analysis decreased LSA PA for RF from 0.25 to 0.21 in FVW. SYK and FVW’s PA did not change between univariate and multivariate analyses; they were nearly identical, with only a minor improvement of 0.01 to 0.02.

**Fig. 4:**
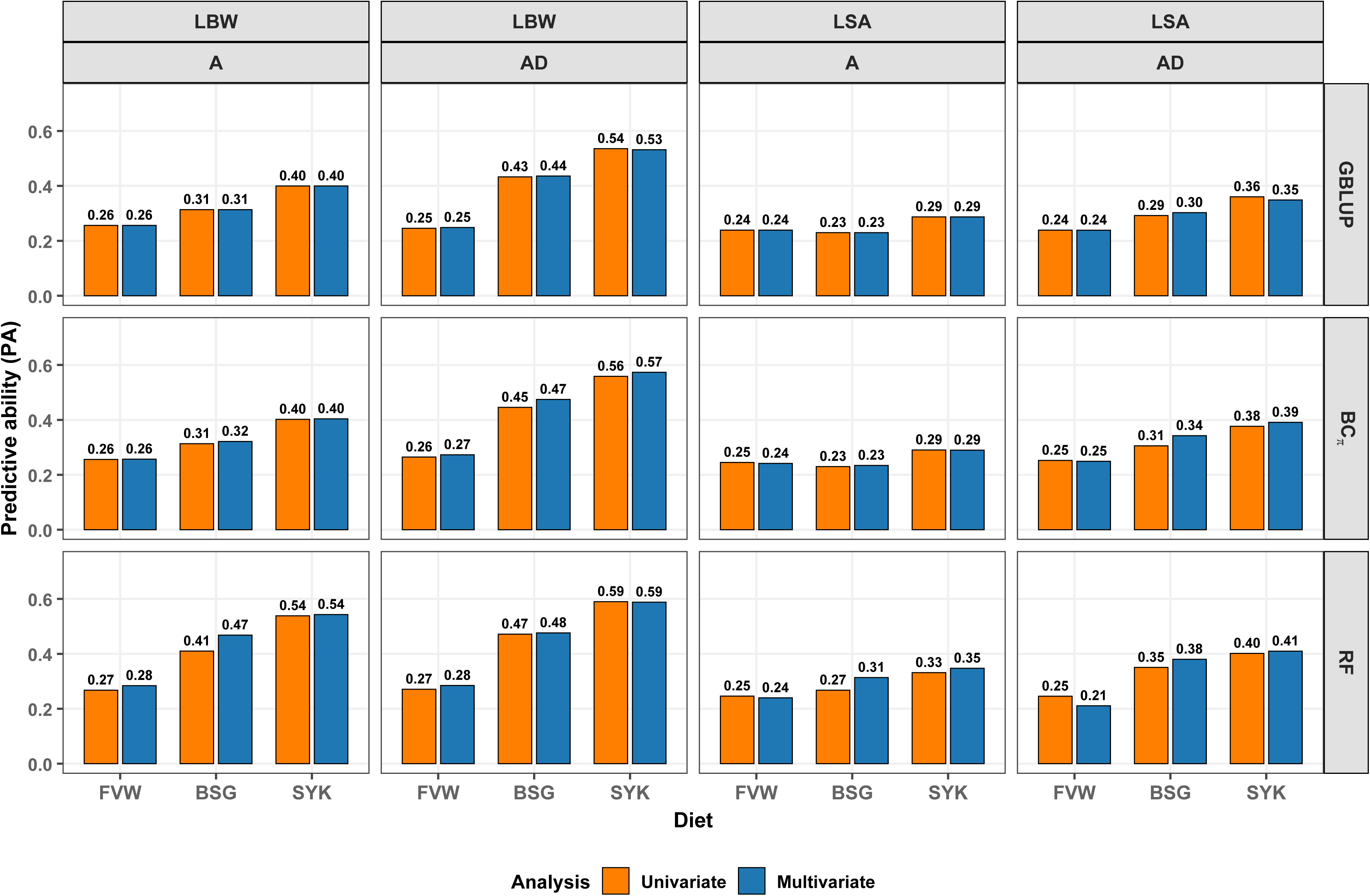
Genomic predictive ability (PA) of different methods: genomic best linear unbiased prediction (GBLUP), Bayes Cπ (BCπ), and random forest (RF), using additive (A) and additive-dominance (AD) models for BSFL growth traits: larval body weight (LBW) and larval surface area (LSA), under univariate and multivariate analyses across diets (SYK: soy-okara, BSG: brewers’ spent grain, FVW: fruit and vegetable waste)

### Cross-diet prediction performance

Cross-diet prediction performance differed across diets, methods, and models (Fig. 5). For both traits (LBW and LSA), training on individuals from SYK and FVW and predicting BSG individuals resulted in the highest PA. In BSG, LBW reached a PA of 0.35 under the A model and 0.40 under the AD model, while LSA reached 0.17 under the A model and 0.31 under the AD model using the RF method. Overall, RF performed best across all the diets and models. However, under the A model for LSA, GBLUP and BCπ showed negative PAs, ranging from 0.03 to −0.02 for SYK and −0.04 for FVW.

**Fig. 5:**
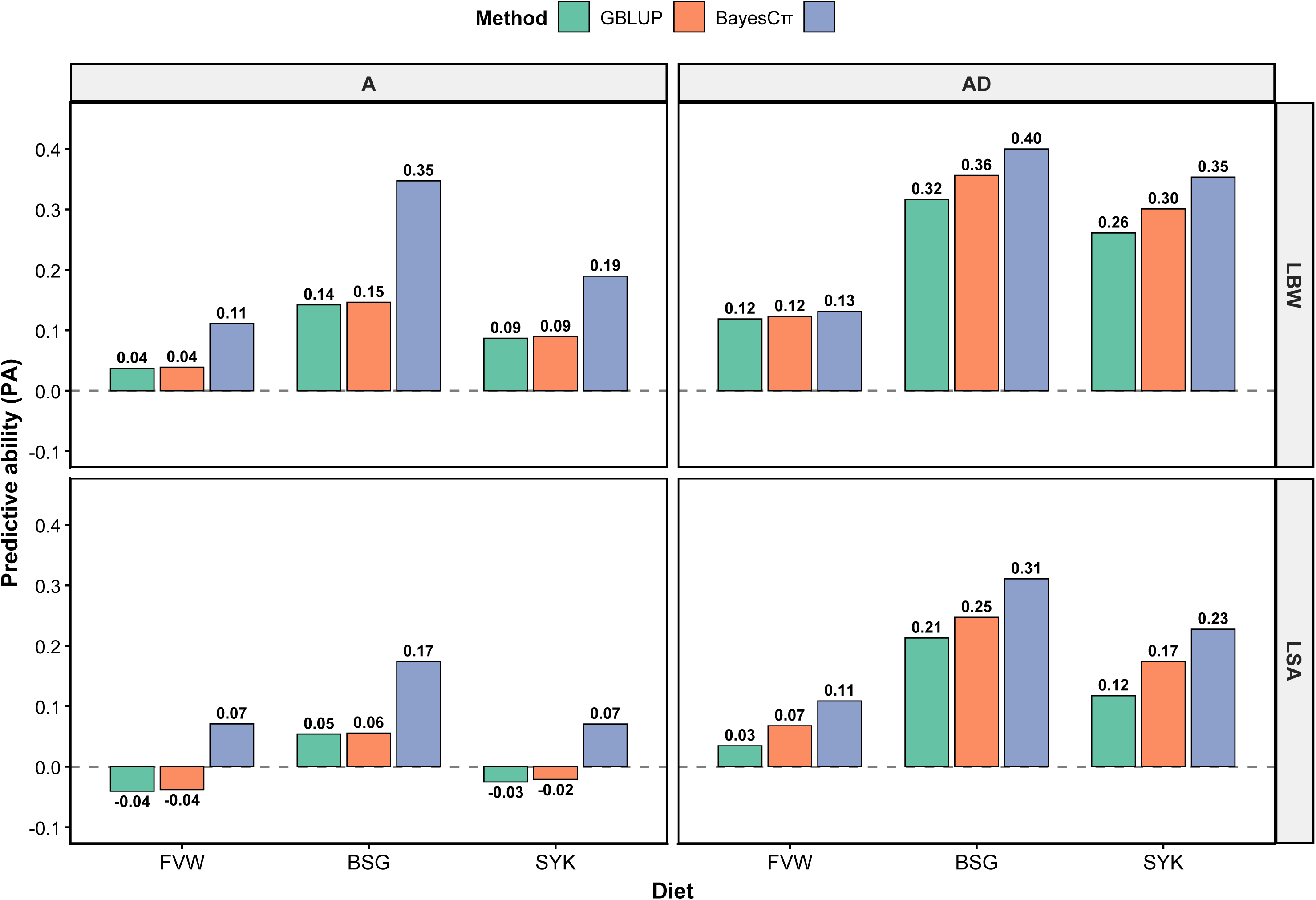
Genomic predictive ability (PA) of different methods: genomic best linear unbiased prediction (GBLUP), Bayes Cπ (BCπ), and random forest (RF), using additive (A) and additive-dominance (AD) models for BSFL growth traits: larval body weight (LBW) and larval surface area (LSA), in cross-diet prediction analyses across diets (SYK: soy-okara, BSG: brewers’ spent grain, FVW: fruit and vegetable waste)

### Effect of SNP density on prediction performance

SNP density influenced PA for both traits (LBW and LSA) across methods and models (Fig. 6). PA increased steadily from 500 to 3,000 SNPs across all evaluation scenarios, then plateaued beyond 3,000 SNPs and remained stable at 4,000 SNPs, with a slight decrease at 4,680 SNPs. AD models consistently achieved higher PA than A models across all densities and methods. In LBW at 3,000 SNPs, AD models reached around 0.40 to 0.44, whereas A models were at 0.29, with the notable exception of RF at 0.41 (Additional file 2, Table S3). The 1,000 top-ranked SNPs yielded higher PA than the full 4,680-SNP panel for both LBW and LSA. However, Bayesian and GBLUP methods performed better than RF under the AD model when using these top 1,000 SNPs (Additional file 1, Figure S4).

**Fig. 6:**
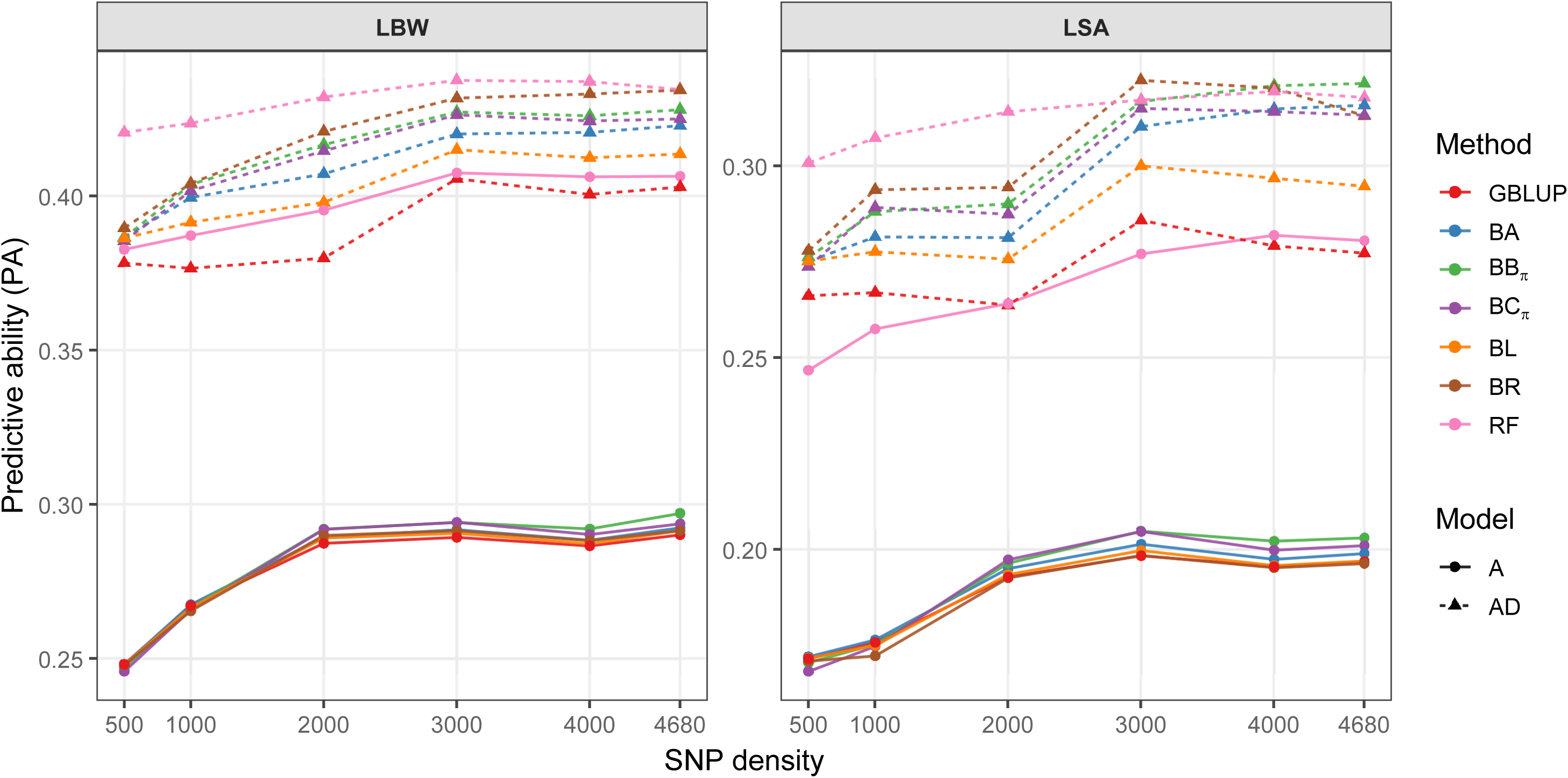
Genomic predictive ability (PA) of different methods: genomic best linear unbiased prediction (GBLUP), Bayes A (BA), Bayes Bπ (BBπ), Bayes Cπ (BCπ), Bayes LASSO (BL), Bayes R (BR), and random forest (RF), using additive (A) and additive-dominance (AD) models across SNP densities for BSFL growth traits: larval body weight (LBW) and larval surface area (LSA)

### Forward and backward prediction performance

Prediction performance differed across the training direction, validation population, traits, diets, methods, and models (Tables 1 and 2). In forward prediction, PA was generally higher under AD models in comparison with A models. Diet-specific predictions were high for SYK, then BSG and FVW, with overall PA reaching 0.24 for both LBW and LSA. In backward prediction, PA was generally lower than in forward prediction, with overall PA reaching 0.19 for LBW and 0.16 for LSA under AD models. Diet-specific PA followed the same trend as forward prediction; SYK achieved higher values, but the BSG showed near zero under the A model, whereas it was positive across all diets in AD models.

**Table 1.** Prediction ability across genomic prediction methods and models for forward prediction of BSFL growth traits in the F7 validation population.

| Trait | Diet | Training (N) | Validation (N) | Model | GBLUP | BCπ | RF |
|---|---|---|---|---|---|---|---|
| LBW | Overall | 2083 | 536 | A | 0.15 | 0.14 | 0.24 |
|  |  |  |  | AD | 0.24 | 0.24 | 0.23 |
|  | SYK | 965 | 536 | A | 0.17 | 0.16 | 0.23 |
|  |  |  |  | AD | 0.22 | 0.24 | 0.22 |
|  | BSG | 555 | 536 | A | -0.04 | -0.01 | 0.19 |
|  |  |  |  | AD | 0.15 | 0.18 | 0.24 |
|  | FVW | 563 | 536 | A | 0.07 | 0.08 | 0.08 |
|  |  |  |  | AD | 0.14 | 0.17 | 0.14 |
| LSA | Overall | 2083 | 536 | A | 0.19 | 0.18 | 0.23 |
|  |  |  |  | AD | 0.23 | 0.24 | 0.24 |
|  | SYK | 965 | 536 | A | 0.21 | 0.21 | 0.16 |
|  |  |  |  | AD | 0.22 | 0.26 | 0.18 |
|  | BSG | 555 | 536 | A | -0.04 | -0.04 | 0.13 |
|  |  |  |  | AD | 0.13 | 0.16 | 0.24 |
|  | FVW | 563 | 536 | A | 0.07 | 0.08 | 0.09 |
|  |  |  |  | AD | 0.11 | 0.13 | 0.20 |

GBLUP, genomic best linear unbiased prediction; BCπ, Bayes Cπ; RF, random forest; A, additive model; AD, additive-dominance model; SYK, soy-okara; BSG, brewers' spent grain; FVW, fruit and vegetable waste; LBW, larval body weight; LSA, larval surface area. Predictions used either the overall base population or diet-specific groups as training populations.

**Table 2.**
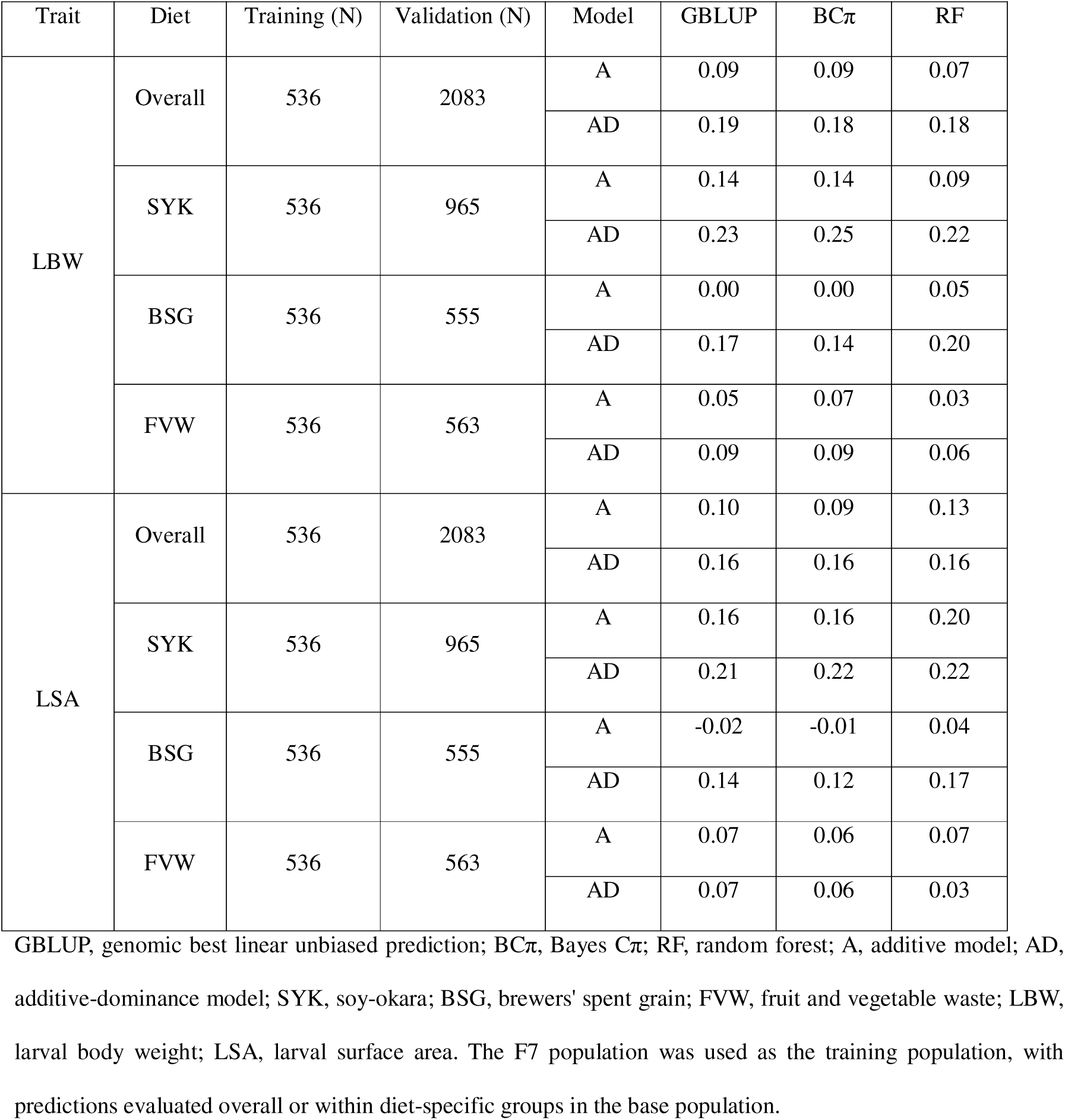
Prediction ability across genomic prediction methods and models for backward prediction of BSFL growth traits in the base population.

| Trait | Diet | Training (N) | Validation (N) | Model | GBLUP | BCπ | RF |
|---|---|---|---|---|---|---|---|
| LBW | Overall | 536 | 2083 | A | 0.09 | 0.09 | 0.07 |
|  |  |  |  | AD | 0.19 | 0.18 | 0.18 |
|  | SYK | 536 | 965 | A | 0.14 | 0.14 | 0.09 |
|  |  |  |  | AD | 0.23 | 0.25 | 0.22 |
|  | BSG | 536 | 555 | A | 0.00 | 0.00 | 0.05 |
|  |  |  |  | AD | 0.17 | 0.14 | 0.20 |
|  | FVW | 536 | 563 | A | 0.05 | 0.07 | 0.03 |
|  |  |  |  | AD | 0.09 | 0.09 | 0.06 |
| LSA | Overall | 536 | 2083 | A | 0.10 | 0.09 | 0.13 |
|  |  |  |  | AD | 0.16 | 0.16 | 0.16 |
|  | SYK | 536 | 965 | A | 0.16 | 0.16 | 0.20 |
|  |  |  |  | AD | 0.21 | 0.22 | 0.22 |
|  | BSG | 536 | 555 | A | -0.02 | -0.01 | 0.04 |
|  |  |  |  | AD | 0.14 | 0.12 | 0.17 |
|  | FVW | 536 | 563 | A | 0.07 | 0.06 | 0.07 |
|  |  |  |  | AD | 0.07 | 0.06 | 0.03 |

GBLUP, genomic best linear unbiased prediction; BCπ, Bayes Cπ; RF, random forest; A, additive model; AD, additive-dominance model; SYK, soy-okara; BSG, brewers' spent grain; FVW, fruit and vegetable waste; LBW, larval body weight; LSA, larval surface area. The F7 population was used as the training population, with predictions evaluated overall or within diet-specific groups in the base population.

## Discussion

This study evaluated genomic prediction approaches for BSFL growth traits, aiming to develop a framework to support genomic selection in BSF and potentially to other insect breeding programmes for food and feed applications. The results indicate that prediction performance varied with trait, nutritional environment, genetic model, marker density, and the genetic relationship between the training and validation populations. These findings establish a foundation to design and evaluate breeding programmes in BSF and future applications.

The prediction performance measured as PA varied among BSFL growth traits across and within diets (Figs. 2 and 3; Additional file 2, Tables S1 and S2); this may reflect differences in trait heritabilities reported in [9] and underlying genetic architecture. LBW showed the highest PA among traits (0.29-0.44), making it an informative trait for genomic prediction, depending on the breeding objective. Prediction performance also varied among statistical methods; GBLUP and Bayesian methods were similar in predictive ability; however, RF reported higher PA for LBW across and within diets. This may be because it captures complex and non-linear relationships among markers compared to the linear GBLUP and Bayesian models, where these relationships are not explicitly modelled [45].

Including dominance in the prediction models improved prediction performance across traits and statistical methods, suggesting dominance contributed to the genomic prediction. However, LBW and LSA in FVW showed no significant difference in PA between A and AD models; this may be because dominance was not significant within FVW. Similarly, accounting for the non-additive genetic effects such as dominance improved prediction performance in sheep [19], chicken and pig [18]. In BSF breeding programmes, founding farm stocks are often sourced from both wild and domesticated populations. Therefore, considering only the additive component may overlook relevant genetic variation in BSFL; both additive and non-additive genetic effects should be evaluated when developing genomic prediction models for BSFL.

The multivariate models which accounted for G × D interactions in the BSFL growth traits performed similarly to or showed marginal improvements in PA compared to the univariate within-diet models. The low genetic correlations reported previously in [9] for BSFL growth traits in different diets may explain the marginal improvement, as information from one diet contributes less to predicting other diets [46]. Accounting for G × D interactions alone doesn’t necessarily improve prediction performance and depends on the magnitude of the genetic correlations.

Cross-diet genomic prediction performance was variable and generally lower than within-diet predictions, depending on the training and target diet combinations. This is consistent with the genetic reranking associated with low genetic correlations among the diets reported in the population [9]. These differences may partly reflect diet-dependent changes in physiological and metabolic pathways, including the expression of genes involved in nutrient digestion and absorption [47]. The reduced performance indicates limited transferability across diets, suggesting models developed for specific diets may not transfer reliably to untested diets. Reference populations should therefore include the major feedstocks of interest, or diet-specific evaluations should be conducted before implementing genomic prediction. For farms that use consistent feedstocks, genomic selection could be tailored to that target diet. Whereas farms that use variable feedstocks need to evaluate the breeding stocks across the diets to develop a line that performs better across the diets or diet-specific ones.

Predictive performance for both LBW and LSA remained stable when marker density was reduced from the full panel (4,680) to 3,000 SNPs across a 1.01 Gb genome, indicating that 3,000 SNPs still retained sufficient marker information and linkage disequilibrium (LD) structure [12]. However, when marker density fell below 3,000 SNPs, prediction performance decreased, possibly because fewer markers were available to adequately capture genomic relationships among individuals and LD structure. This suggests that 3,000 SNPs may be sufficient for this population to maintain prediction performance, potentially reducing genotyping costs. The LD patterns reported in [12] provide information for the selection of informative markers and the development of cost-effective, low-density panels for BSF breeding in Australian populations. The shared ancestry of Australian BSF following colonisation may have contributed to the observed LD patterns. The top-ranked 1,000 SNPs from GBLUP performed better than the full panel, indicating that the prioritised SNPs retain substantial predictive information, even with fewer markers. Parametric methods (GBLUP and Bayesian) outperformed RF on this panel, which may reflect the advantage of selecting markers from GBLUP. Recent studies indicate that routine breeding programmes in livestock, plant and aquaculture are starting to use low-cost panels of 3 to 10K SNPs [30, 31, 48, 49]. However, further evaluation using larger, more diverse populations across generations and economic evaluation are needed to establish whether reduced panels can maintain performance while reducing the cost.

Prediction performance declined across multiple generations, consistent with previous studies in honey bee [50], dairy cattle [51], spruce [52], and shrimp [53]. This decline may reflect changes in genetic relationships between the training and validation populations across generations, as well as changes in allele frequency and quantitative trait loci as a result of recombination. As genetic relationships between the reference population and selection candidates decline, genomic information estimated in the original population also becomes less informative in subsequent generations. Additionally, differences in the production environments between the base and F7 populations, including diet, rearing conditions and locations, may have contributed to the reduction in the prediction performance through genotype-by-environment interactions. The *F_ST_* of 0.063 ± 0.004 between the base and F7 populations also supports genetic differentiation and is consistent with changes in population structure across generations. These results highlight the need to periodically update the reference population and account for production environment differences in training models to obtain accurate estimates in this short-generation-interval species.

Genomic prediction has also been evaluated in other insect species such as the honey bee, where prediction accuracy ranged from 0.23 to 0.65 for honey yield and workability traits [50]. Our results extend the application of genomic prediction to BSF growth traits by accounting for both additive and dominance effects. However, its uptake in insect breeding remains in its infancy compared to aquaculture and livestock breeding [2, 17, 31]. Nonetheless, application of genomic selection in aquaculture species with similar life-history traits to insects can provide valuable insights into applying genomic prediction to commercial insect production systems [17, 27, 30, 31]. For example, genomic selection studies in shrimp demonstrate the importance of carefully managing family contributions and genetic relatedness between reference and target populations when developing genomic prediction strategies in high fecundity species with vulnerable life stages and short generation intervals [53]. These biological similarities offer established strategies for managing large family contributions and traceability. However, their implementation needs to account for the BSF’s biology and production systems.

This study showcased the potential of genomic prediction in BSFL; however, its implementation will depend on breeding population structure and management of genetic diversity. Maintaining equal family contributions across life stages or in selection across generations could help manage genetic diversity. However, its influence on genetic gain and prediction performance needs to be evaluated. The decline in predictive ability across generations necessitates evaluating strategies for maintaining prediction performance, including updating the reference population to account for connectivity changes between generations. The effect of mate choice, including non-random mating and contributions, on genetic gain also needs to be evaluated. Furthermore, transferability of genomic prediction to the external germplasm remains to be established, particularly given the reduced predictive ability observed in forward and backward prediction scenarios. This study did not investigate unrelated populations; further research should evaluate the prediction performance in genetically differentiated populations. Future research also needs to investigate whether genomic predictions can support selection of breeding lines under commercial conditions rather than at the individual level.

Accurate and efficient phenotypic strategies are required for BSF breeding programmes, with computer vision approaches being investigated as a high-throughput potential [54]. Because of the short generation interval, rapid, non-invasive DNA extraction from larvae/adults is necessary for genotyping. However, it is not yet standardised, as one study used adult legs for DNA extraction [55]. Genotyping must also be economical compared with the additional genetic gain achieved through genomic selection. Therefore, researchers need to develop rapid, standardised, cost-effective workflows. These findings should be interpreted in the context of the population, diets, and the limited number of generations evaluated. Importantly, the realised genetic gain from genomic selection remains to be evaluated.

Based on the findings of this study, the following recommendations are proposed for genomic prediction in BSF breeding programmes:

• Genetic diversity management: Maintain balanced family contribution across life stages, sampling and selection to manage genetic diversity and inbreeding;

• Genomic prediction reference population: Establish a sufficiently large reference population that represents the target rearing environments, and maintain genetic connectedness with the selection population;

• Mating strategies: Evaluate mate allocation and non-random mating designs to manage diversity, control inbreeding and improve genetic gain;

• G × D interactions: Evaluate and account for G × D interactions when developing generalist or diet-specific breeding programmes;

• Genetic variance and prediction model: Evaluate additive and non-additive genetic variance and appropriately account for both components in the prediction models. Regularly update and evaluate prediction models for their transferability across generations, genetically different populations and external germplasm;

• Breeding line selection: Extend and evaluate genomic prediction for estimating genetic merit at the breeding line or strain-level under commercial production conditions;

• Cost-effective genotyping: Design and validate cost-effective rapid, low-density genotyping strategies suitable for BSF production systems; and

• Commercial evaluation: Evaluate realised genetic gain and the cost-benefit of genomic prediction under commercial conditions.

## Conclusions

Our study demonstrates the potential of genomic prediction in BSFL growth traits and provides a framework for BSF breeding programmes. Prediction performance differed by trait, diet, model, statistical method, SNP density, and generation. Accounting for dominance improved prediction, and accounting for G × D added limited improvement because genetic correlations among diets were low. Cross-diet and cross-generation prediction reduced transferability, highlighting the importance of representing major feedstocks and breeding generations in the reference populations. Periodic evaluation and updating of the reference population may help maintain the prediction performance. Prediction performance remained stable at the 3,000-SNP panel, highlighting the low-cost genotyping. Implementing genomic prediction needs standardised high-throughput phenotyping, rapid genotyping, and standardised breeding evaluation and workflow in commercial breeding conditions.

## Declarations

Ethics approval and consent to participate

## Not applicable

Consent for publication

Not applicable.

## Availability of data and materials

The phenotypic and SNP genotypic data supporting this study will be made available after publication; currently, they are embargoed and archived in the Research Data JCU repository: https://doi.org/10.25903/t0bn-a463. The scripts used for the analyses are available at https://github.com/KB-Gowda/BSF_Quantitative_Genomics.git.

## Authors’ contributions

KBG and KRZ conceived and designed the study. KBG curated data, performed the analysis, and wrote the first draft. KBG, CM, DRJ and KRZ contributed to reviewing and revising the manuscript. DRJ and KRZ secured the funding, and KRZ provided the project administration, resources and supervision. All authors read and approved the final manuscript.

## Acknowledgements

Authors acknowledge the BSF research teams at JCU and FlyFarm Australia for their assistance with the experiments and data collection.

## Additional files

### Additional file 1: Figure S1

Format: figure in PDF file

Title: Distributions of overall adjusted phenotypes (residuals) for larval body weight (LBW), larval length (LL), larval width (LW), and larval surface area (LSA)

### Additional file 1: Figure S2

Format: figure in PDF file

Title: Distributions of diet-wise adjusted phenotypes (residuals) for larval body weight (LBW) and larval surface area (LSA) across three diet groups: Fruit and vegetable waste (FVW), Brewer’s spent grain (BSG), and Soy-okara (SYK)

### Additional file 1: Figure S3

Format: figure in PDF file

Title: Genome-wide marker density distribution across seven chromosomes (Chr1-Chr7) evaluated for six SNP panel densities (4,680, 4,000, 3,000, 2,000, 1,000, and 500 SNPs). Marker density is measured in 1 Mb physical windows along each chromosome

### Additional file 1: Figure S4

Format: figure in PDF file

Title: Genomic predictive ability (PA) of different methods: genomic best linear unbiased prediction (GBLUP), Bayes A (BA), Bayes Bπ (BBπ), Bayes Cπ (BCπ), Bayes LASSO (BL), Bayes R (BR), and random forest (RF), using additive (A) and additive-dominance (AD) models across top (1,000 SNPs) and full panels (4,680 SNPs) for BSFL growth traits: larval body weight (LBW) and larval surface area (LSA)

### Additional file 2: Table S1

Format: table in Excel file

Title: Predictive ability (± standard deviation), bias (± standard deviation), and mean squared error of prediction (MSEP ± standard deviation) across genomic prediction methods: genomic best linear unbiased prediction (GBLUP), Bayes A (BA), Bayes Bπ (BBπ), Bayes Cπ (BCπ), Bayes Lasso (BL), Bayes R (BR), and random forest (RF), using additive (A) and additive-dominance (AD) models for four BSFL growth traits: larval body weight (LBW), larval length (LL), larval width (LW) and larval surface area (LSA)

### Additional file 2: Table S2

Format: table in Excel file

Title: Predictive ability (± standard deviation), bias (± standard deviation), and mean squared error of prediction (MSEP ± standard deviation) across genomic prediction methods: genomic best linear unbiased prediction (GBLUP), Bayes A (BA), Bayes Bπ (BBπ), Bayes Cπ (BCπ), Bayes Lasso (BL), Bayes R (BR), and random forest (RF), using additive (A) and additive-dominance (AD) models for within-diet (SYK: soy-okara, BSG: brewers’ spent grain, FVW: fruit and vegetable waste) prediction of BSFL growth traits: larval body weight (LBW) and larval surface area (LSA)

### Additional file 2: Table S3

Format: table in Excel file

Title: Predictive ability (± standard deviation), bias (± standard deviation), and mean squared error of prediction (MSEP ± standard deviation) across genomic prediction methods: genomic best linear unbiased prediction (GBLUP), Bayes A (BA), Bayes Bπ (BBπ), Bayes Cπ (BCπ), Bayes Lasso (BL), Bayes R (BR), and random forest (RF), using additive (A) and additive-dominance (AD) models across SNP densities for BSFL growth traits: larval body weight (LBW) and larval surface area (LSA)

