## Supplementary figures for "A genomic prediction framework for growth traits in black soldier fly (*Hermetia illucens*) breeding programmes"

### Additional file 1

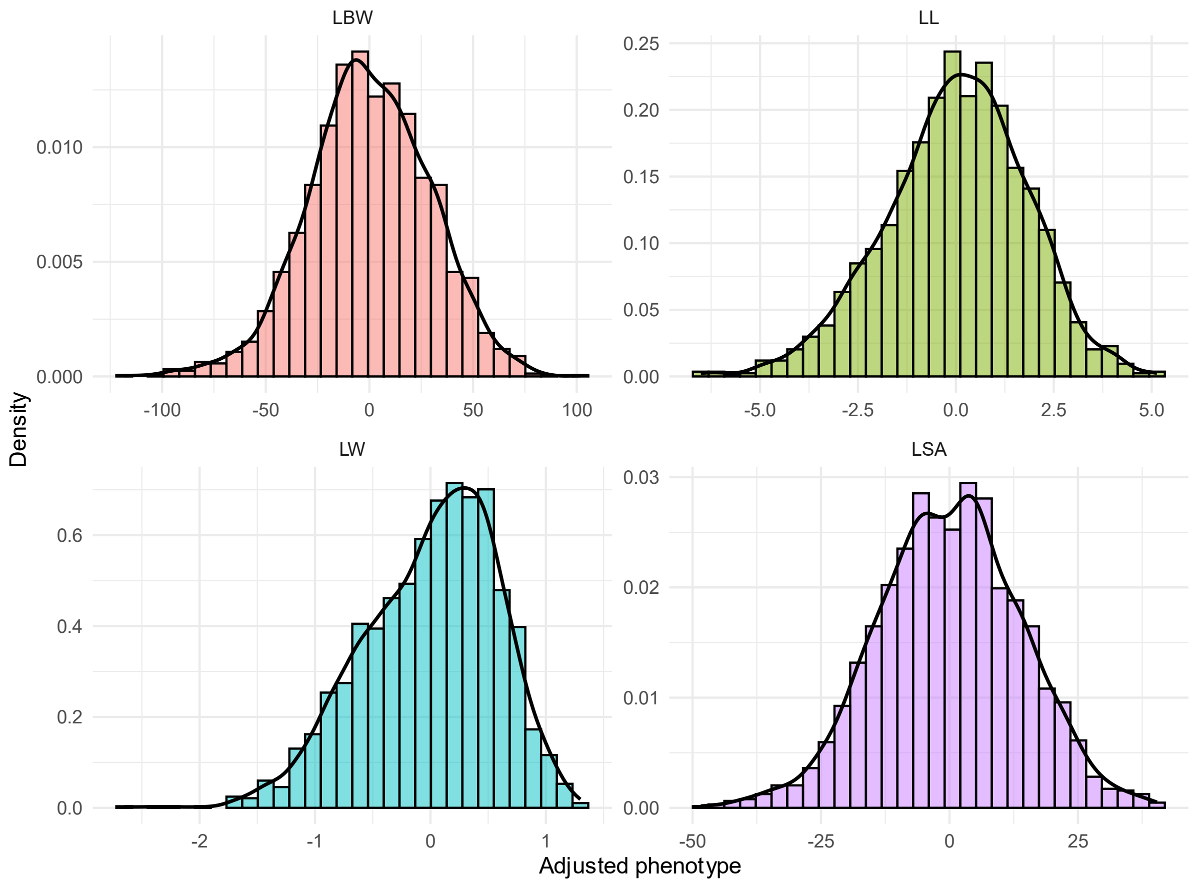


**Figure S1**. Distributions of overall adjusted phenotypes (residuals) for larval body weight (LBW), larval length (LL), larval width (LW), and larval surface area (LSA)

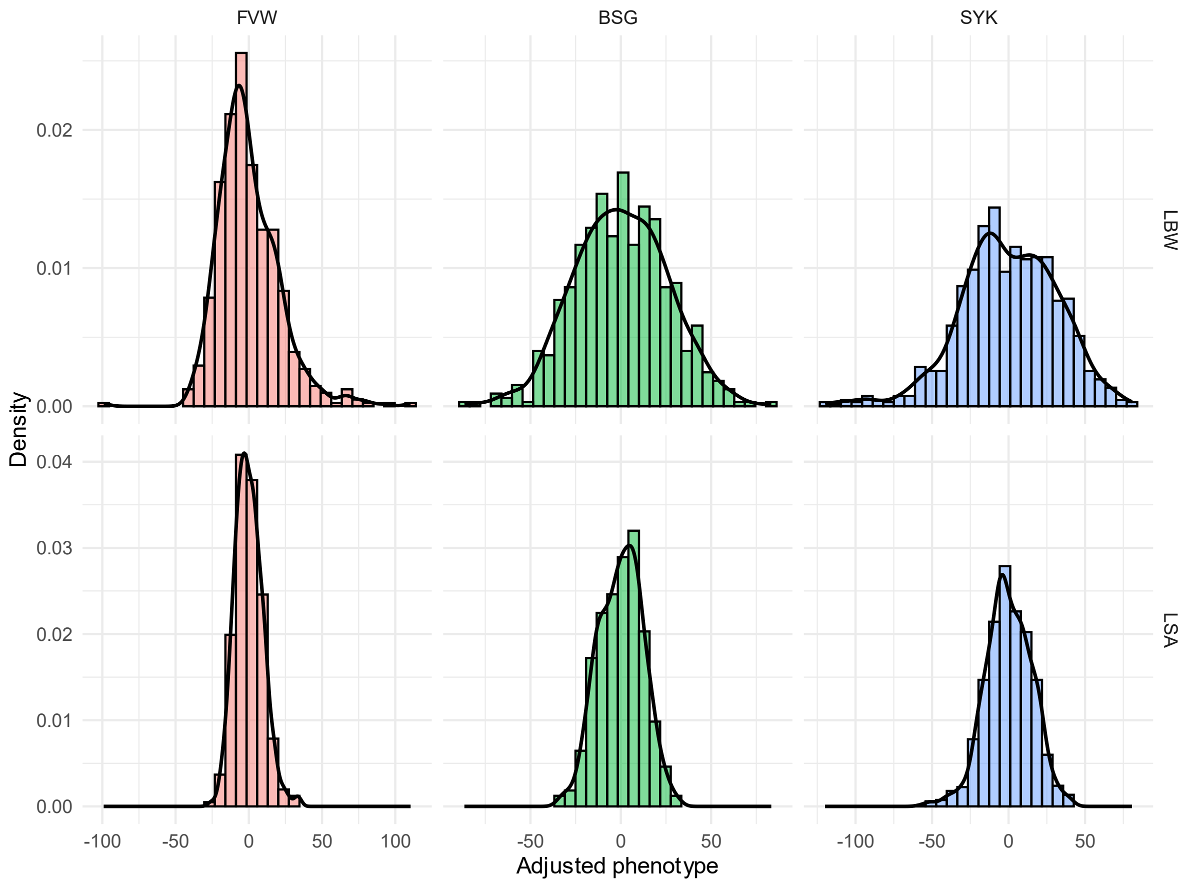


**Figure S2**. Distributions of diet-wise adjusted phenotypes (residuals) for larval body weight (LBW) and larval surface area (LSA) across three diet groups: Fruit and vegetable waste (FVW), Brewer's spent grain (BSG), and Soy-okara (SYK)

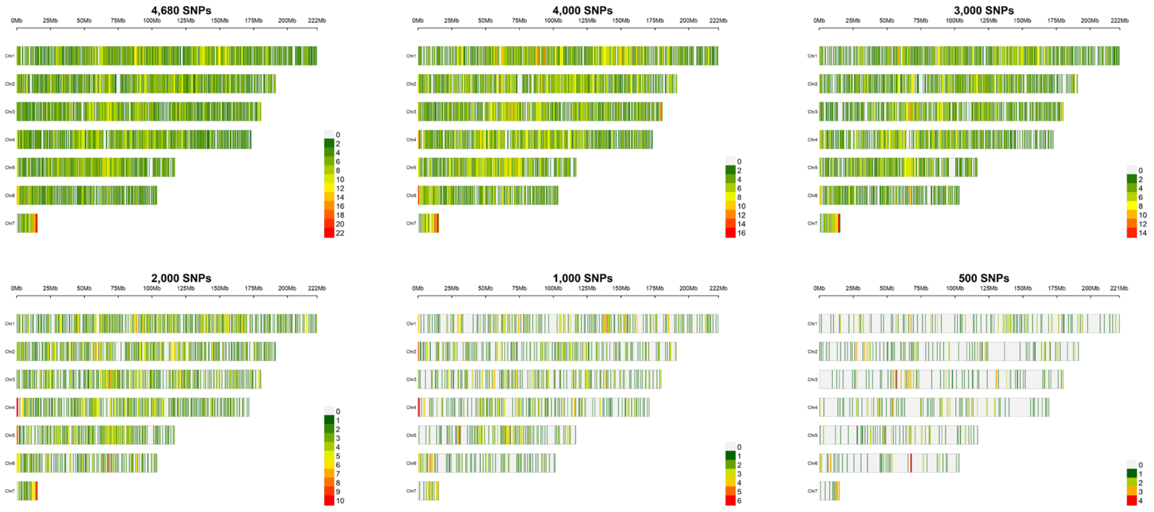


**Figure S3**. Genome-wide marker density distribution across seven chromosomes (Chr1-Chr7) evaluated for six SNP panel densities (4,680, 4,000, 3,000, 2,000, 1,000, and 500 SNPs). Marker density is measured in 1 Mb physical windows along each chromosome

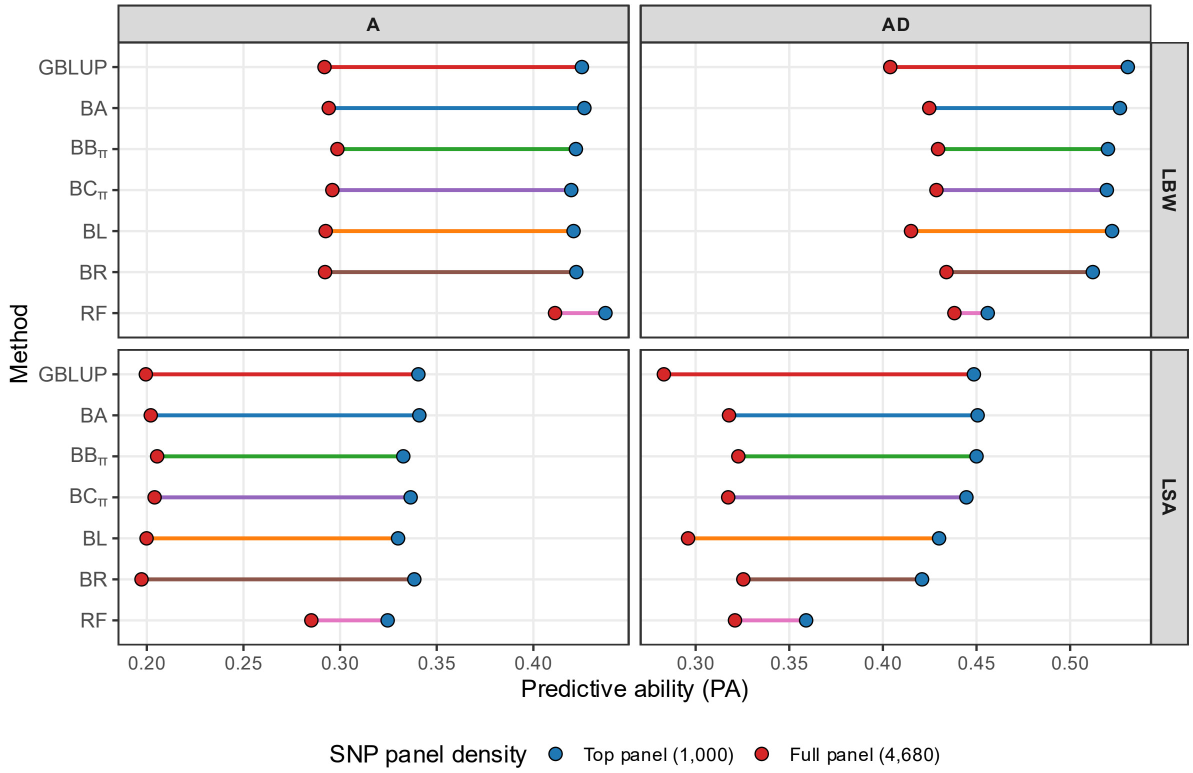


**Figure S4**. Genomic predictive ability (PA) of different methods: genomic best linear unbiased prediction (GBLUP), Bayes A (BA), Bayes Bπ (BBπ), Bayes Cπ (BCπ), Bayes LASSO (BL), Bayes R (BR), and random forest (RF), using additive (A) and additive-dominance (AD) models across top (1,000 SNPs) and full panels (4,680 SNPs) for BSFL growth traits: larval body weight (LBW) and larval surface area (LSA)
